# YTHDF3 promotes the progression of gastric cancer by activating the Wnt/β-catenin signaling pathway by targeting NEK7

**DOI:** 10.1101/2025.01.25.634861

**Authors:** Xiaoran Sun, Yifan Chen, Kunyi Liu, Yi Si, Mingda Xuan, Rui Zhang, Jiao Jiao, Ting Liu, Shuangshuang Han, Jia Wang, Weifang Yu

**Affiliations:** Gastrointestinal Disease Diagnosis and Treatment Center, The First Hospital of Hebei Medical University, Shijiazhuang, China; Gastroenterology Department, The First Affiliated Hospital of Hebei North University, Zhangjiakou, China; Department of Infectious Diseases, The First Hospital of Hebei Medical University, Shijiazhuang, China

**Keywords:** Gastric cancer, YTHDF3, NEK7, Wnt/β-catenin

## Abstract

Gastric cancer (GC) is a widespread and deadly malignancy among global digestive tract tumors, alarmingly ranked the fifth most common cancer in terms of incidence and mortality. This study used immunohistochemistry and RT-qPCR to reveal a significant upregulation of YTHDF3 expression in GC tissues compared with that of adjacent normal tissues. This increased expression of YTHDF3 was closely associated with the incidence of lymph node metastasis and the progression to more advanced TNM stages. *In vitro* experiments demonstrated that YTHDF3 was crucial in promoting the aggressive characteristics of GC cells by enhancing their ability to proliferate, migrate, and invade, while simultaneously inhibiting apoptosis. The role of YTHDF3 in the progression of GC was further confirmed using the BALB/c nude mice subcutaneous tumor model, which showed a slowed tumor growth after YTHDF3 knockdown. This study further confirmed through Co-IF RIP-qPCR, western blotting, and rescue studies that YTHDF3 promotes GC progression by targeting NEK7 to regulate the Wnt/β-catenin signaling pathway. In conclusion, our pivotal findings highlight the YTHDF3/NEK7/Wnt/β-catenin axis as a promising and accessible therapeutic target for the development of new treatments aimed at improving clinical outcome of patients with GC.

## Introduction

Gastric cancer (GC) is one of the most common malignant tumors of the digestive tract worldwide. In 2022, the incidence and mortality rates of GC ranked fifth among all cancers globally, accounting for 4.9% of new cancer cases and 6.8% of cancer deaths (1). The occurrence and progression of GC may involve the interaction between oncogenes and tumor suppressor genes, genetics, and environmental factors, and it is a complex multi-step process. Although early-stage patients with GC achieve a good cure rate and prognosis, advanced GC often has a poor prognosis and high mortality rate due to the lack of timely and effective treatment methods. With the establishment of public biological databases for GC and the development of bioinformatics, an increasing number of molecules are considered to be associated with the progression of GC. Therefore, the in-depth study of the molecular mechanisms underlying the development and progression of GC plays a crucial role in the early diagnosis and precise treatment of GC.

The YTH domain family proteins (YTHDFs), including YTHDF1, YTHDF2, and YTHDF3, are a class of cytoplasmic m6A-binding proteins defined by the vertebrate YTH domain, which play a broad role in regulating the fate of RNA. The YTHDF family mediates tumor proliferation, metastasis, and metabolism and has the potential to serve as predictive and therapeutic biomarkers. The levels of YTH domain proteins are elevated in many malignant tumors (2–4). Studies indicate that members of the YTH domain protein family, YTHDF1, YTHDF2, and YTHDF3, play a crucial role in the regulation of RNA fate and are closely associated with tumor proliferation, metastasis, and metabolism. This phenomenon is observed in various types of cancer, including hepatocellular carcinoma, prostate cancer, pancreatic cancer, GC, lung cancer, and colorectal cancer (5–10).

As a significant member of the YTHDF family, YTHDF3, as other YTHDF members, contains a C-terminal YTH domain and a N-terminal domain rich in P/Q/N residues. YTHDF3 is more complex and context-dependent. Its precise molecular function, key downstream effectors, and the mechanistic basis for its action remain largely unexplored. Recent studies have shown that YTHDF3 can directly interact with YTHDF2, affecting mRNA degradation. It can also interact with YTHDF1 to improve mRNA translation (11), suggesting its potential to modulate diverse signaling pathways and perform multiple roles *in vivo*. Emerging evidence implicates YTHDF3 in the regulation of cancer progression in various tumor types. In triple-negative breast cancer, YTHDF3 stabilizes ZEB1 mRNA, promoting progression and metastasis (12). Similarly, in ocular melanoma, it improves cancer stem cell tumorigenicity by promoting CTNNB1 translation (13) and in osteosarcoma, it facilitates aerobic glycolysis by stabilizing PGK1 (14). YTHDF3 can promote the degradation of GAS5 mRNA, thus enhancing the progression and invasive potential of colorectal cancer (15). In pancreatic cancer, YTHDF3 engages in a negative feedback loop that affects glycolysis and tumorigenesis (16), whereas in hepatocellular carcinoma, it promotes progression through the YTHDF3/m6A-EGFR/STAT3 axis and EMT (17). Additionally, high expression of YTHDF3 in colorectal cancer is associated with poor pathological differentiation, advanced stage, and metastasis, driving cancer cell proliferation and migration (18). These findings underscore the complex and context-dependent roles of YTHDF3 in cancer development, making it a promising target for therapeutic intervention.

However, the exact role of YTHDF3 in the occurrence and progression of GC remains unclear. We found that YTHDF3 is highly expressed in GC and is closely associated with the poor prognosis of patients with GC. YTHDF3 enhances the malignant phenotype of GC cells and promotes the growth of GC *in vivo*. Through transcriptome sequencing, bioinformatics analysis, Co-IF/RIP-qPCR experiments, and rescue studies, we first discovered that the downstream target gene for YTHDF3 in GC is NEK7, which is also highly expressed in GC and interacts with YTHDF3. YTHDF3 upregulates the expression of the NEK7 protein and activates the Wnt/β-catenin signaling pathway, further enhancing the proliferation, migration, and invasion capabilities of GC cells, ultimately promoting the progression of GC.

## Materials and Methods

### Patients and samples

We acquired 50 pairs of GC tissues and adjacent normal tissues from the Hebei Medical University’s First Hospital. Before surgery, none of the patients had undergone chemotherapy or radiation, and all had signed informed consent forms. Following isolation, the aforementioned tissue samples were immediately placed in liquid nitrogen before being moved to the First Hospital’s Biological Sample Bank at Hebei Medical University for storage. Pathological confirmation of gastric adenocarcinoma/signet-ring cell carcinoma was obtained for the cancer tissues used in this investigation. This study complies with the principles of the Declaration of Helsinki and was authorized by the Ethics Committee of the First Hospital of Hebei Medical University (Approval letter No. S00996).

### Cell culture

Wuhan Pricella Biotechnology Co., Ltd. provided the human gastric mucosa cell (GES-1) and GC cell lines (HGC-27, AGS, MKN7, MKN74), which were confirmed by STR analysis. Cells were grown in RPMI 1640 medium (Gibco, Gaithersburg, MD, USA) with 1% penicillin-streptomycin solution (Solarbio Sciences & Technology Co. Ltd, Beijing, China) and 10% fetal bovine serum (FBS; Gibco) added as supplements. Cells were maintained in an incubator with 5% carbon dioxide at 37°C and a steady temperature and humidity level. The Wnt/β-catenin pathway inhibitor, IWR-1-endo, was obtained from Selleck Chemicals (Houston, TX, USA). PCR was used to detect mycoplasma monthly. The cell lines were utilized for a maximum of 25 passages in each experiment.

### Transfections, stable cell lines, siRNAs, shRNAs, and plasmids

Lipofectamine 3000 (Invitrogen, Carlsbad, CA, USA) was used for transfection when cell confluence in a cell culture flask or plate reached 80–90%. HGC-27. AGS cells were transfected with YTHDF3 siRNA and a negative control (GenePharma Co., Ltd., Shanghai, China), whereas MKN7 and MKN74 cells were transfected with a YTHDF3 overexpression plasmid and a negative control (GenePharma). HGC-27 cells were also transfected with YTHDF3 shRNA and a negative control (GenePharma). MKN74 cells were transfected with NEK7 siRNA and a negative control (GenePharma), whereas HGC-27 cells were transfected with a NEK7 overexpression plasmid and a negative control (GenePharma). Experiments were conducted 48 hours following transfection. For YTHDF3 shRNA transfection in HGC-27 cells, the cell culture medium was modified to add neomycin at a final concentration of 600 μg/mL (without other antibiotics) 48 hours following transfection to establish stable transfected cell lines (HGC-27 shNC, HGC-27 shYTHDF3). Following the formation of the clones, the neomycin concentration was raised to 300 μg/mL and cultured overnight.

### RNA extraction and RT-qPCR

Using the RNA-Easy Isolation Reagent (Vazyme Biotech Co., Ltd., Nanjing, China), total RNA was extracted from cells and tissues. The PrimeScript RT reagent kit (Takara, Beijing, China) was then used to reverse-transcribe (RT) the RNA into complementary DNA (cDNA). The AceQ Universal SYBR qPCR Master Mix (Vazyme) was used for RT-qPCR, and the endogenous control mRNA was β-actin. Every sample was examined three times. The CT (cyclic threshold) value for each sample was used to assess the mRNA expression level, and the 2^-ΔΔCT^ method was used to quantify the relative expression level of each sample.

### Immunohistochemistry

Tissue samples obtained from patients and naked mice were preserved in 4% paraformaldehyde, then sectioned, stained, and paraffin embedded (Shangha YiYang Instrument Co., Ltd., Shanghai, China). Immunohistochemistry (IHC) analysis was performed using the Rabbit/Mouse two-step detection kit (Beijing ZSBG-Bio Co., Ltd., Beijing, China) in accordance with the manufacturer’s instructions. IHC staining was performed using antibodies against YTHDF3 (diluted 1:500; Proteintech, Wuhan, Hubei, PRC.; Catalog number: 25537-1-AP), Ki67 (diluted 1:100; Proteintech; Catalog number: 27309-1-AP), and NEK7 (diluted 1:50; Santa Cruz Biotechnology, CA, USA; Catalog number: sc-393539). The average optical density (AOD) and immunoreactive score (IRS) method were used to examine the staining results.

### Western blotting

A protease/phosphatase inhibitor (Solarbio) and RIPA lysis buffer (Solarbio) were used in a 100:1 ratio to extract the protein. The proteins were transferred to a polyvinylidene fluoride membrane (Merck Millipore, Billerica, MA, USA) after these were electrophoretically separated on a 10% SDS polyacrylamide gel (Bio-Rad Laboratories Inc., Hercules, CA, USA). In this study, the following antibodies were used: β-actin antibody (diluted 1:1500; ZSBG-Bio; Catalog number: TA-09), YTHDF3 antibody (diluted 1:1000; Proteintech, Wuhan, Hubei, PRC; Catalog number: 25537-1-AP), Cyclin D1 antibody (diluted 1:1000; Proteintech; Catalog number: 60186-1-lg), c-MYC antibody (diluted 1:500; Proteintech; Catalog number: 10828-1AP), β-catenin antibody (diluted 1:1000; Proteintech; Catalog number: 51067-2-AP). Phospho-Beta Catenin Recombinant monoclonal antibody(diluted 1:1000; Proteintech; Catalog number: 80067-1-RR). The Odyssey Scanning System (LI-COR Biosciences, Lincoln, NE, USA) identified the immunoreactive protein bands.

### Cell proliferation assay

The cells for each experimental group were evenly spaced in 96-well plates at a density of 2×10^3^ cells per well and cultured in full culture medium for the CCK-8 test. To each well 10 μL of the Cell Counting Kit -8 (CCK-8; Dojindo, Tokyo, Japan) reagent was added at 24, 48, 72, and 96 hours after distribution. After that, the cells were kept in the incubator for two more hours. The Promega GloMax luminescence detector (Promega, Madison, WI, USA) was used to measure the absorbance at 450 nm of each well.

For the colony formation test, the cells in each group were evenly distributed at a density of 1000 cells per well in 6-well plates. Every three days, a new culture medium was added. After culture for 7 to 14 days, the cells were washed twice with PBS, treated for 30 minutes with 4% paraformaldehyde, then stained for 20 minutes with 0.1% crystal violet.

### Cell migration assay

The cells of each group were evenly distributed in 6-well plates for the wound healing assay. A 200 μL pipette tip was used to draw two straight lines in each well to represent the wound once the cell confluence was reached or was approximately 100%. Following two PBS washes, the cells were cultivated in a culture medium devoid of FBS. Following the creation of the wound, each well was photographed at 0 and 48 hours. The migration rate was calculated as the ratio of the gap width at 0 to 48 hours.

A total of 4×10^4^ (HGC-27, AGS) and 8×10^4^ (MKN7, MKN74) cells were introduced into the upper chamber (Corning Incorporated, Corning, NY, USA) in 200 μL FBS-free media for the Transwell migration test. A volume of 700 μL of complete medium was introduced into the lower chamber to induce downward migration. Cells on the upper chamber side of the polycarbonate membrane were extracted with a swab after a 48-hour culture period. After two PBS washes, the chamber was treated for 30 minutes with 4% paraformaldehyde and then for 20 minutes with 0.1% crystal violet. PBS was used once more to clean the excess crystal violet. ImageJ software (National Institutes of Health, Bethesda, MD, USA) was used to count the stained cells after five visual areas were chosen at random for photography.

### Cell invasion assay

To promote downward invasion of cells in the upper chamber, cells were resuspended in 100 μL of FBS-free medium and introduced to the upper chamber. Meanwhile, 600 μL of complete medium was added to the lower chamber for the Transwell invasion assay. All other experimental protocols were identical to the Transwell migration assay protocol.

### Cell apoptosis assay

Fort-eight hours after transfection, cells were extracted from each treatment group and stained with the Annexin V-FITC/PI apoptosis assay kit (NeoBioscience, Shenzhen, China). 1×10^5^ cells were subjected to apoptosis analysis using flow cytometry (BD Biosciences, San Jose, CA, USA). The percentage of apoptotic cells was determined using the FlowJo software.

### TUNEL Assay

Cell apoptosis was detected by the TUNEL (Terminal deoxynucleotidyl transferase-mediated dUTP Nick-End Labeling) assay using a commercial kit by following the manufacturer’s instructions (Beyotime Biotechnology, Shanghai, China; Catalog number: C1090). Briefly, AGS and HGC-27 cells were seeded on sterile coverslips placed in 6-well plates. After cell confluence had reached 50% to 70%, the cells were gently rinsed once with PBS. The cells were then fixed with 4% paraformaldehyde for 30 minutes at room temperature and permeabilized with 0.3% Triton X-100 (Solarbio; Catalog number: T8200) for 10 minutes. Following two washes with PBS, the cells were incubated with 50 μL of TUNEL detection solution per sample for 60 minutes at 37°C in the dark. After three washes with PBS, cell nuclei were counterstained with 4’,6-diamidino-2-phenylindole (DAPI; Beyotime Biotechnology) for 10 minutes. The coverslips were then mounted onto glass slides, and representative images were captured using a laser scanning confocal microscope (ZEISS, Oberkochen, Germany).

### Animal studies

Ten male BALB/c nude mice weighing 16 to 20 g and 4 to 6 weeks of age were purchased from Beijing Huafukang Biotechnology Co., LTD. They were randomly split into two groups, with five mice in each group, and housed in a pathogen-free environment with sufficient food and water. The room temperature was kept at 22°C with a 12-/12-hour light/dark cycle. After disinfecting the skin of the left anterior axillary fossa of the nude mice, a sterile syringe was used to subcutaneously inject HGC-27 cells (5×10^6^ cells per mouse) that had been stably transfected with shRNA-NC and shRNA-YTHDF3. Tumor size was measured every 2 days beginning on day 3 after injection using the formula volume = (long diameter × short diameter 2)/2. The nude mice were euthanized when the tumor volume reached 1000 mm^3^. The excised tumor tissues were sectioned and embedded in paraffin for HE and IHC examination. The aforementioned studies were conducted in accordance with the experimental animal care and use system and authorized by the First Hospital’s Ethics Committee at Hebei Medical University.

### RNA sequencing

After 24 hours of siRNA-NC and siRNA-YTHDF3 transfection, total RNA was collected from HGC-27 cells. Beijing Novogene Technology Co., LTD was assigned the task of RNA sequencing (RNA seq) by creating the RNA library and performing the sequencing analysis. Briefly, 6 samples (siNC × 3 and siYTHDF3 × 3) were examined using Illumina’s NovaSeq 6000 platform. To obtain high-quality data, each sample’s output data was subjected to quality control and filtering. High-quality data were aligned to reference genomes using the HISAT software, which was also used to measure the sample’s levels of gene or transcript expression. Differential genes were classified as those with log2 |Fold Change| >0 and *P<*0.05. DESeq2 software was utilized for determining differentially expressed genes (DEG) investigation.

### Co-Immunofluorescent assay

In a 6-well plate, HGC-27 cells were seeded uniformly on coverslips. Cells were gently washed three times with PBS once the cell fusion rate had reached 20% to 30%. For the Co-Immunofluorescent (Co-IF) assay, the cells were first blocked with 2% bovine serum albumin (BSA-V; Solarbio; Catalog number: A8020), fixed with 4% paraformaldehyde, and permeabilized with 0.2% Triton X-10 (Solarbio; Catalog number: T8200). YTHDF3 antibody (diluted 1:50; Proteintech; Catalog number: 25537-1-AP) and NEK7 antibody (diluted 1:50; Santa Cruz Biotechnology; Catalog number: sc-393539) were then added to the cells and incubated overnight at 4°C. The cells were then incubated for 1 hour at room temperature in the dark in the presence of fluorescent secondary 309 antibody (diluted 1:500; Cy3 anti-rabbit IgG, FITC anti-mouse IgG; Beyotime Biotechnology Co., Shanghai, China).Lastly, the nuclei were stained with 4’6’-diamino-2-phenylindole dihydrochloride (DAPI; Beyotime Biotechnology Co.). A laser scanning confocal microscope (ZEISS, Oberkochen, Germany) was used to take representative pictures.

### RIP-qPCR

In accordance with the instructions of the RNA Immunoprecipitation Kit (BersinBio, Guangzhou, China; Catalog number: Bes5101), HGC-27 cells were collected and lysed. An YTHDF3 specific antibody (10 µg; Proteintech; Catalog number: 25537-1-AP) was used to bind the endogenous expression of the YTHDF3 protein in cells, precipitate the target protein-RNA complex, and then isolate and purify the RNA within the complex, RNA was then subjected to quantitative real-time PCR (qPCR) analysis.

### RNA decay assay

Actinomycin D (ACTD) was introduced into the HGC-27 cell culture at 1 μg/mL. RNA was extracted at 0, 1, 2, and 4 hours for RT-qPCR. Gene expression was normalized to NEK7 mRNA levels at 0 h.

### Immunofluorescence assay

HGC-27 cells (1×10^4^ cells/mL) were cultured on 1 cm glass slides. After fixation with 4% paraformaldehyde (100,496; Sigma-Aldrich) at room temperature, the cells were permeabilized and blocked with 0.5% Triton X-100 (83,443; Sigma-Aldrich). The cells were subsequently incubated with a β-catenin antibody (1:50, 51067-2-AP, Proteintech) and secondary antibodies (cy3 1:500, Beyotime Biotechnology, Shanghai, China; Catalog number: A0516). The nuclei were treated with 4,6-diamino-2-phenyl indole (DAPI, Beyotime Biotechnology). Images were acquired using a laser scanning confocal microscope (ZEISS, Oberkochen, Germany). Image J software (version 1.8.0, National Institutes of Health) was used to circle the nuclear area and analyze the mean fluorescence intensity in a blinded fashion.

### Bioinformatics analysis

TCGA (https://www.aclbi.com/static/index.html#/tcga) and GEO (https://www.aclbi.com/static/index.html#/geo) databases were interrogated to determine the expression of YTHDF3 mRNA in cancer and surrounding tissues of patients with GC. Gene expression levels in GC cases were examined using GEPIA2 (http://gepia2.cancer-pku.cn/#index), together with nearby normal sublocalization within cells. Primers were constructed using PrimerBank (https://pga.mgh.harvard.edu/primerbank/) and the Primer designing tool (https://www.ncbi.nlm.nih.gov/tools/primer-blast/). BioRender (https://www.biorender.com/) was used to create schematic diagrams.

### Statistical analysis

For each group of experiments, three independent replicates were conducted. Data with a normal distribution were displayed as mean ± standard deviation, whereas data with a non-normal distribution were displayed as median and interquartile distance. SPSS v.26.0 (IBM, Armonk, NY, USA) and GraphPad Prism v.9.5 (GraphPad Software, La Jolla, CA, USA) were used to perform the statistical analysis. The Student’s t-test, one-way analysis of variance (ANOVA), and two-way ANOVA were among the statistical techniques used. A P-value < 0.05 was deemed statistically significant.

### Data Availability

The corresponding author can provide the data produced in this study upon request. The Cancer Genome Atlas (TCGA) and Gene Expression Omnibus (GEO) datasets GSE54129 and GSE66229 were used in this analysis.

## Results

### YTHDF3 was upregulated in patients with GC and was associated with a poor prognosis

To investigate the role of YTHDF3 in the pathophysiology of GC, we first analyzed its expression in GC and adjacent normal tissues using TCGA database, as well as the GSE54129 and GSE66229 datasets. Our analysis revealed a significant upregulation of YTHDF3 expression in GC tissues compared with adjacent normal tissues (Figure 1A-C). This observation was further supported by the higher expression of YTHDF3 in tissues of stomach adenocarcinoma (STAD) as assessed using the online tool GEPIA2 (Figure 1D). Given the limited understanding of the function of YTHDF3 in GC, we selected it as the target gene for this study. We quantified YTHDF3 mRNA expression in 12 pairs of fresh-frozen GC tissues and adjacent normal tissues using RT-qPCR. The results demonstrated a statistically significant increase in YTHDF3 mRNA expression in GC tissues (*P<*0.01), consistent with the TCGA and GEO data (Figure 1E). To validate these findings at the protein level, we performed IHC on the GC and adjacent normal tissue. The IHC results revealed that the YTHDF3 protein was predominantly localized in the cytoplasm of GC cells and was significantly overexpressed in GC tissues compared with adjacent normal tissues (*P<*0.001, x2 = 12.19) (Figure 1F, Table 1). Subsequently, we explored the relationship between YTHDF3 expression and the clinical characteristics of patients with GC. YTHDF3 expression was found to be independent of sex, age, tumor size, and tumor differentiation. However, it was significantly associated with lymph node metastasis (*P<*0.01) and TNM staging (*P<*0.05) (Table 2). These data suggest that YTHDF3 overexpression is associated with a poor prognosis in patients with GC, highlighting its potential as a significant biomarker and therapeutic target in GC.

**Figure 1.**
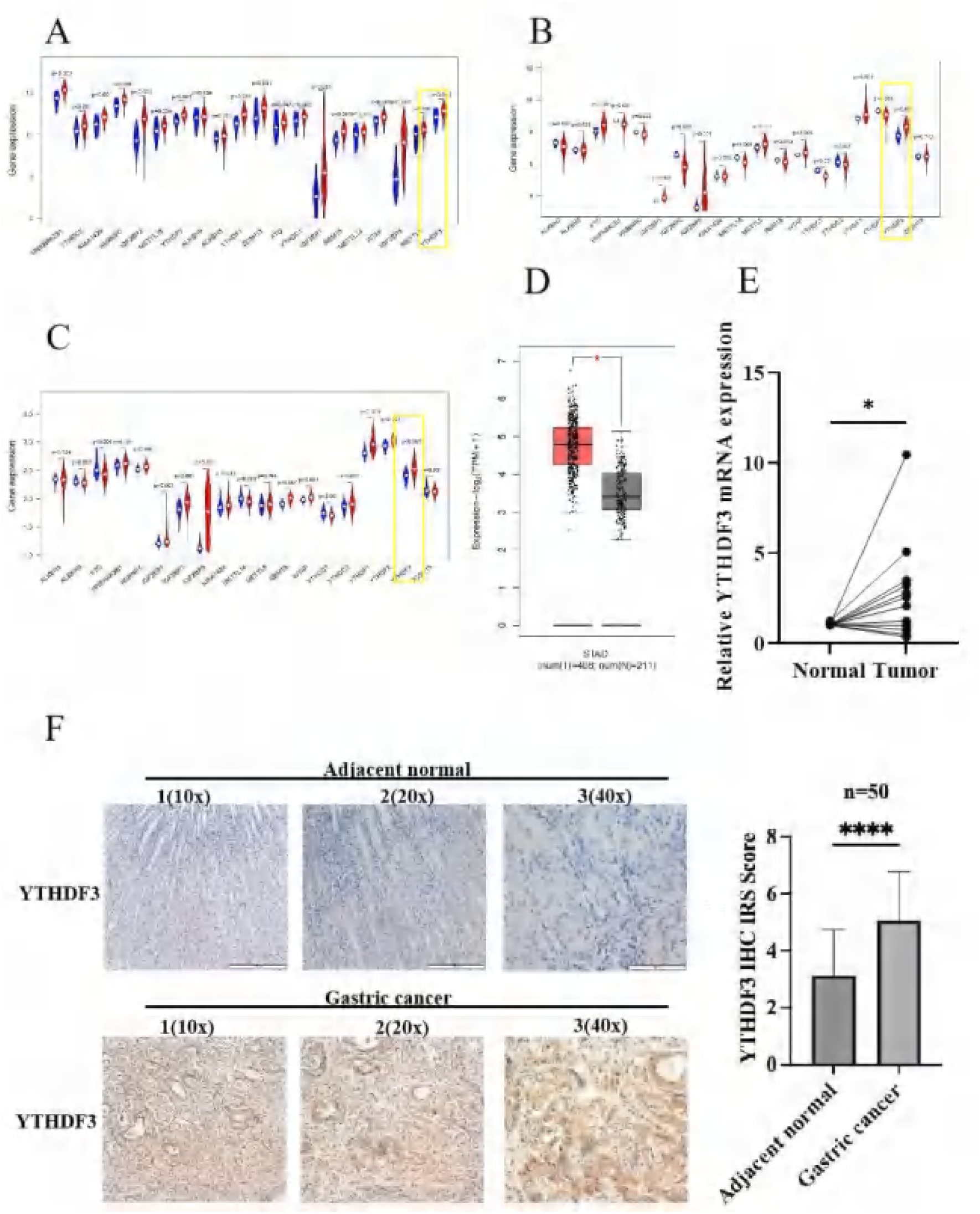
Increased YTHDF3 expression is associated with a poor prognosis in patients with GC. (A-C) Expression of YTHDF3 in TCGA database, and in the GSE54129 and GSE66229 datasets. (D) YTHDF3 is highly expressed in STAD tissues based on the analysis of the online databases GEPIA2. (E) YTHDF3 mRNA expression in 12 pairs of GC and adjacent normal tissue samples as detected by RT-qPCR. (F) Representative images showing the expression of YTHDF3 protein in 50 pairs of GC and adjacent normal tissue samples, as detected by IHC. Scale bar, 20 μm. Data are presented as means ± SD. \**P<*0.05

**Table 1.**
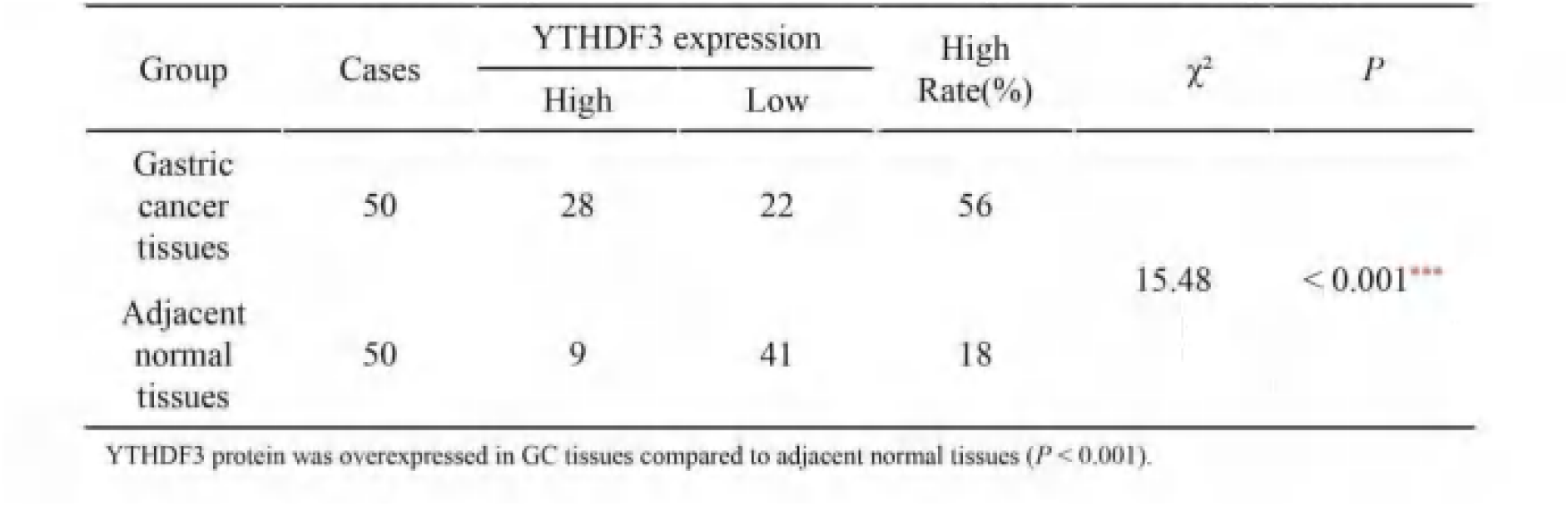
The expression level of YTHDF3 protein in two groups.

| Group | Cases | YTHDF3 expression | | High Rate(%) | $\chi^2$ | *P* |
| --- | --- | --- | --- | --- | --- | --- |
|  |  | High | Low |  |  |  |
| Gastric cancer tissues | 50 | 28 | 22 | 56 | 15.48 | < 0.001*** |
| Adjacent normal tissues | 50 | 9 | 41 | 18 |  |  |

YTHDF3 protein was overexpressed in GC tissues compared to adjacent normal tissues ( $P < 0.001$ ).

**Table 2.** The relationship between the expression of YTIIDF3 in gastric cancer tissue and the clinicoapathological data of patients.

| Group | Cases | YTHDF3 expression | | $\chi^2$ | *P* |
| --- | --- | --- | --- | --- | --- |
|  |  | High | Low |  |  |
| Gender |  |  |  |  |  |
| Male | 37 | 20 | 17 | 0.219 | 0.64 |
| Female | 13 | 8 | 5 |  |  |
| Age (years) |  |  |  |  |  |
| ≤ 60 | 11 | 5 | 6 | 0.636 | 0.425 |
| > 60 | 39 | 23 | 16 |  |  |
| Tumor size (cm) |  |  |  |  |  |
| ≤ 4 cm | 27 | 15 | 12 | 0.005 | 0.945 |
| > 4 cm | 23 | 13 | 10 |  |  |
| Differentiation |  |  |  |  |  |
| Poorly | 26 | 14 | 12 | 0.102 | 0.749 |
| Well | 24 | 14 | 10 |  |  |
| Lymphnode metastasis |  |  |  |  |  |
| Absent | 16 | 5 | 11 | 5.850 | 0.016** |
| Present | 34 | 23 | 11 |  |  |
| TNM Staging |  |  |  |  |  |
| I + II | 21 | 8 | 13 | 4.711 | 0.030* |
| III + IV | 29 | 20 | 9 |  |  |

YTHDF3 expression was associated with lymph node metastasis ( $P < 0.05$ ) and TNM staging ( $P < 0.05$ ), but not with gender, age, tumor size, or tumor differentiation.

### Knockdown of YTHDF3 suppressed proliferation, migration, and invasion and promoted the apoptosis of GC cells *in vitro*

We first downregulated YTHDF3 expression in HGC-27 and AGS cell lines and examined the impact on several cellular processes to investigate the mechanisms of YTHDF3 activity in GC. A considerable decrease was observed in the levels of YTHDF3 mRNA (Figure 2A, B) and protein (Figure 2C, D) after transfection with siRNA targeting YTHDF3 (siYTHDF3) and a siRNA negative control (siNC). The CCK-8 assay demonstrated that cell viability in HGC-27 and AGS cells was significantly reduced following YTHDF3 knockdown when compared with the control condition (Figure 2E, F). The wound healing capacity of YTHDF3 knockdown cells was significantly altered compared with that of controls in HGC-27 and AGS cells after 48 hours (Figure 2G, H). The number of apoptotic cells in HGC-27 and AGS cell lines increased when YTHDF3 expression was silenced, as indicated by flow cytometry analysis (Figure 2I, J). The number of apoptotic cells in AGS cell lines increased when YTHDF3 expression was silenced, as indicated by the Tunel assay (Figure 2K). Furthermore, the results of the Transwell assay indicated a significant reduction in both the number of migrated and invasive cells in the YTHDF3 knockdown group for both HGC-27 and AGS cell lines (Figure 2L–O). These findings imply that downregulation of YTHDF3 stimulates apoptosis and suppresses invasion, migration, and proliferation of GC cells.

**Figure 2.**
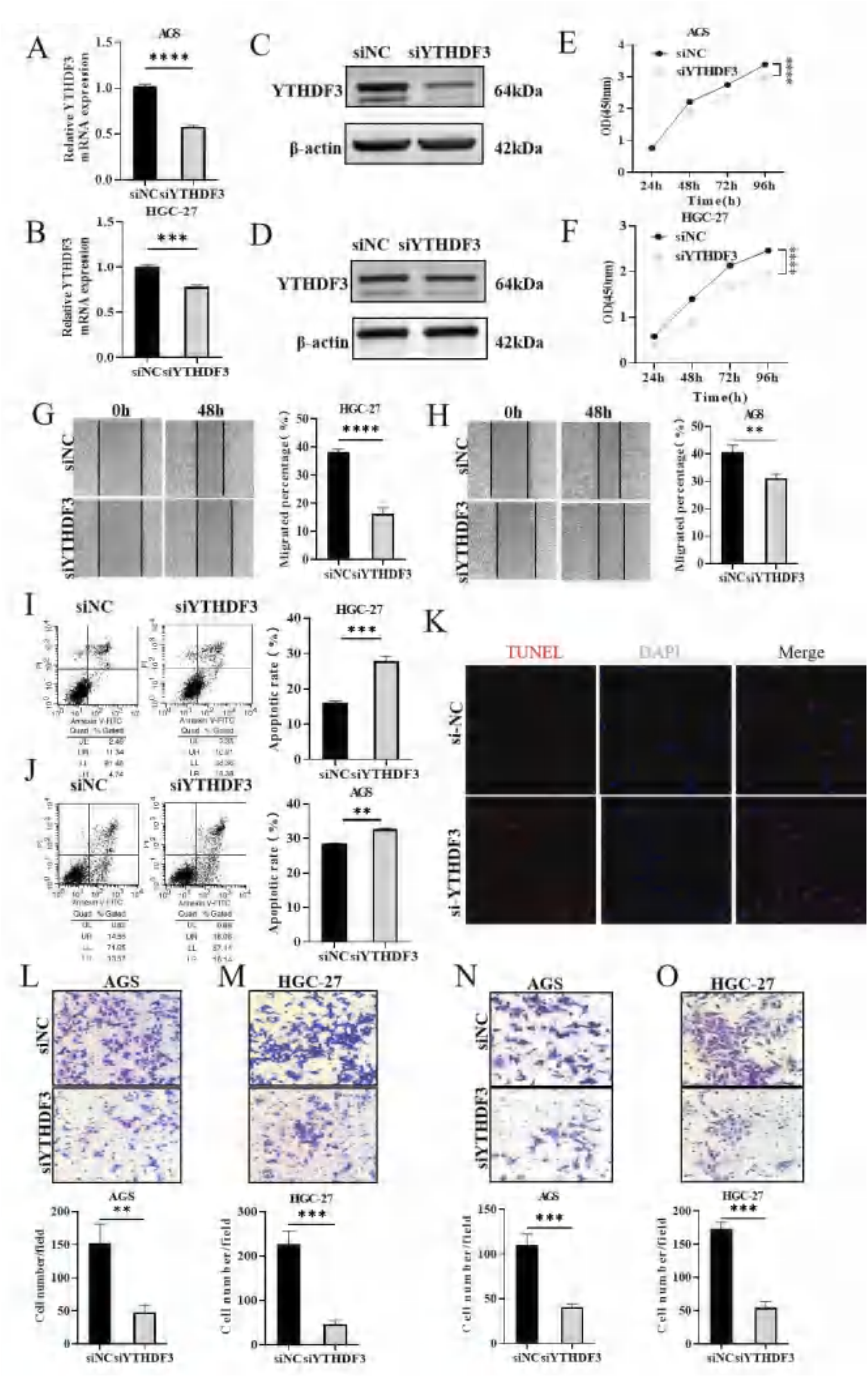
Knockdown of YTHDF3 in GC cells suppresses cell proliferation, migration, and invasion *in vitro*. (A, B) YTHDF3 mRNA expression in HGC-27 and AGS cells as detected by RT-qPCR. (C, D) YTHDF3 protein expression in HGC-27 and AGS as detected by WB. (E, F) Effects of YTHDF3 knockdown on the proliferation of HGC-27 and AGS cells as determined by the CCK-8 assay. (G, H) Effect of YTHDF3 knockdown on HGC-27 and AGS cell migration was assessed using a wound healing assay. Scale bar, 200 μm. (I, J) The effect of YTHDF3 knockdown on apoptosis of HGC-27 and AGS cells as determined by flow cytometry. (K) Effects of YTHDF3 knockdown on the apoptosis of AGS cells as determined by the Tunel assay. (L, M) Effects of YTHDF3 knockdown on HGC-27 and AGS cell migration as evaluated using the Transwell migration assay. Scale bar, 100 μm. (N, O) Effects of YTHDF3 knockdown on HGC-27 and AGS cell invasion as assessed using a Transwell invasion assay. Scale bar, 100 μm. Data are presented as means ± SD. \**P<*0.05, \*\**P<*0.01, \*\*\**P<*0.001, \*\*\*\**P<*0.0001

### YTHDF3 overexpression promoted proliferation, migration, and invasion and suppressed apoptosis of GC cells *in vitro*

We upregulated YTHDF3 expression in MKN7 and MKN74 cell lines to observe its impact on cellular biological functions. Following transfection of cells with pcDNA3.1-YTHDF3 (YTHDF3) and pcDNA3.1-vector (Vector), we observed a marked increase in mRNA levels of YTHDF3 (Figure 3A, B) and protein levels (Figure 3C, D). The CCK-8 assay revealed a significant improvement in cell viability in MKN7 and MKN74 cells following overexpression of YTHDF3, compared with the control group (Figure 3E, F). The wound healing ability of cells with upregulation of YTHDF3 was significantly improved compared with the control group in MKN7 and MKN74 cells after 48 hours (Figure 3G, H). The number of apoptotic cells in the MKN7 and MKN74 cell lines decreased when YTHDF3 expression was elevated, according to the flow cytometry study (Figure 3I, J). The number of apoptotic cells in MKN74 cell lines decreased when YTHDF3 expression was elevated, according to the Tunel assay (Figure 3K). Furthermore, the results of the Transwell assay demonstrated a significant increase in both the number of migrated and invasive cells in the upregulated group of YTHDF3 for the MKN7 and MKN74 cell lines (Figure 3L–O). altogether, these findings imply that YTHDF3 may function as a pro-oncogene in GC, as overexpression of the protein can increase invasion, migration, and proliferation of GC cells while preventing apoptosis.

**Figure 3.**
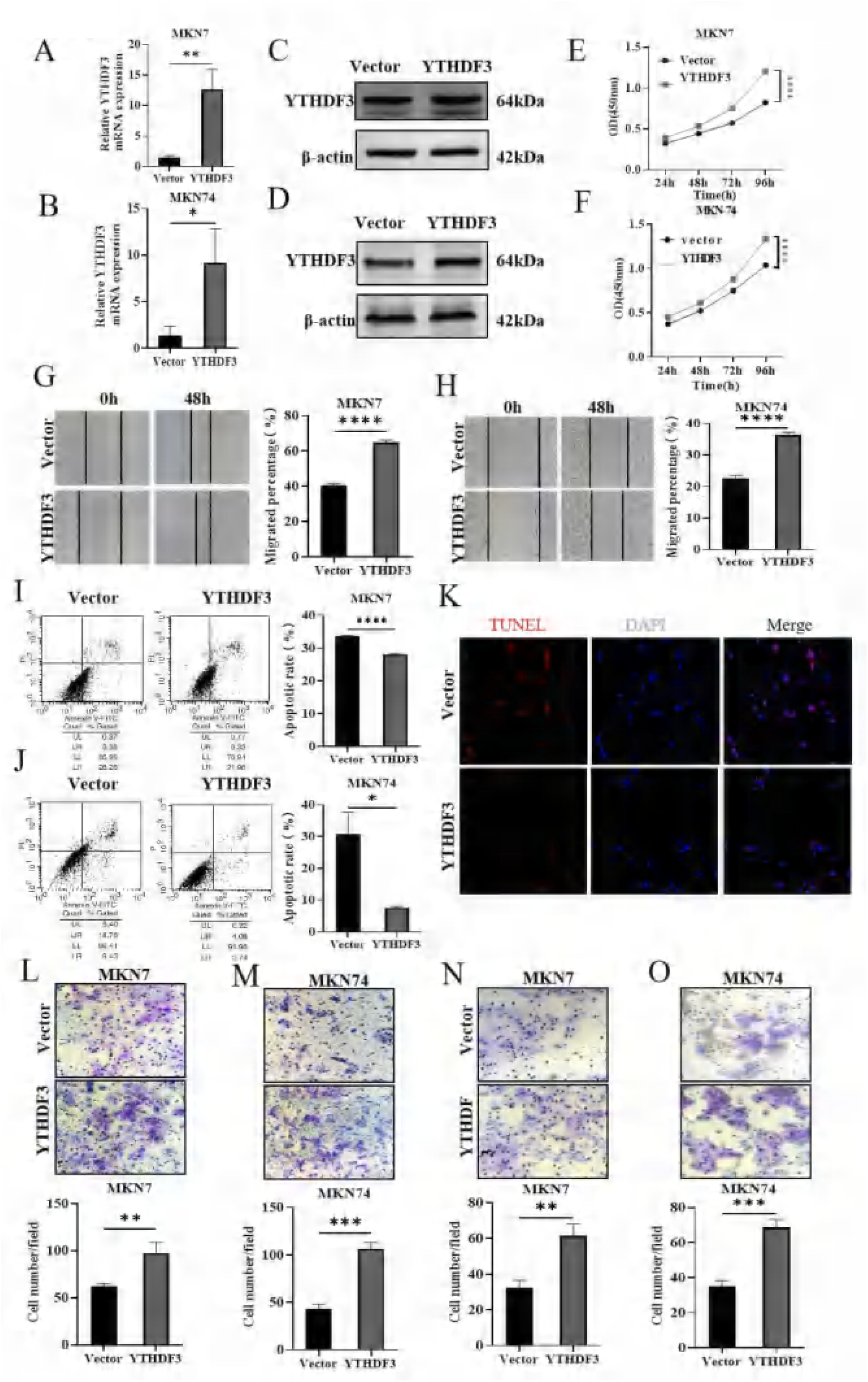
Overexpression of YTHDF3 in GC cells and human gastric mucosa cells promotes cell proliferation, migration, and invasion *in vitro*. (A, B) The expression of YTHDF3 mRNA in MKN7 and MKN74 as detected by RT-qPCR. (C, D) Expression of the YTHDF3 protein in MKN7 and MKN74 as detected by western blotting. (E, F) Effects of YTHDF3 overexpression on MKN7 and MKN74 cell proliferation were assessed using the CCK-8 assay. (G, H) Effect of YTHDF3 overexpression on MKN7 and MKN74 cell migration as evaluated by a wound healing assay. Scale bar, 200 μm.(I, J) Effects of YTHDF3 overexpression on MKN7 and MKN74 cell apoptosis as analyzed by flow cytometry. (K) Effects of YTHDF3 overexpression on apoptosis of MKN74 cells as analyzed using the Tunel assay. (L, M) Effects of YTHDF3 overexpression on MKN7 and MKN74 cell migration as determined using a Transwell migration assay. Scale bar, 100 μm. (N, O) Effects of YTHDF3 overexpression on MKN7 and MKN74 cell invasion as assessed by a Transwell invasion assay. Scale bar, 100 μm. Data are presented as means ± SD. \*\**P<*0.01, \*\*\**P<*0.001, \*\*\*\**P<*0.0001

### YTHDF3 deficiency inhibited the growth of GC *in vivo*

We created tumor-bearing nude mouse models with reduced expression of YTHDF3 and equivalent negative controls to better understand the mechanism of YTHDF3 in GC *in vivo*. After successfully establishing HGC-27 cell lines that were stably transfected with shRNA-YTHDF3 and shRNA-negative control (shNC/shYTHDF3), cells were subcutaneously injected into two groups of nude mice, respectively. Figure 4A–D shows the tumor-bearing models as well as the tumor volume curves, actual tumors, and tumor weights. The growth rate, volume, and weight of the tumors in the shYTHDF3 group were significantly lower than those of the shNC group. The morphological characteristics of the xenograft tissues were assessed by HE staining, and the findings showed that the cell structure and morphology resembled those of the GC tissues (Figure 4E). Compared with the shNC group, the proliferation capacity of cells in the shYTHDF3 group was greatly suppressed, according to Ki67 immunohistochemical staining (Figure 4F). These results imply that GC growth can be inhibited in vivo by YTHDF3 knockdown.

**Figure 4.**
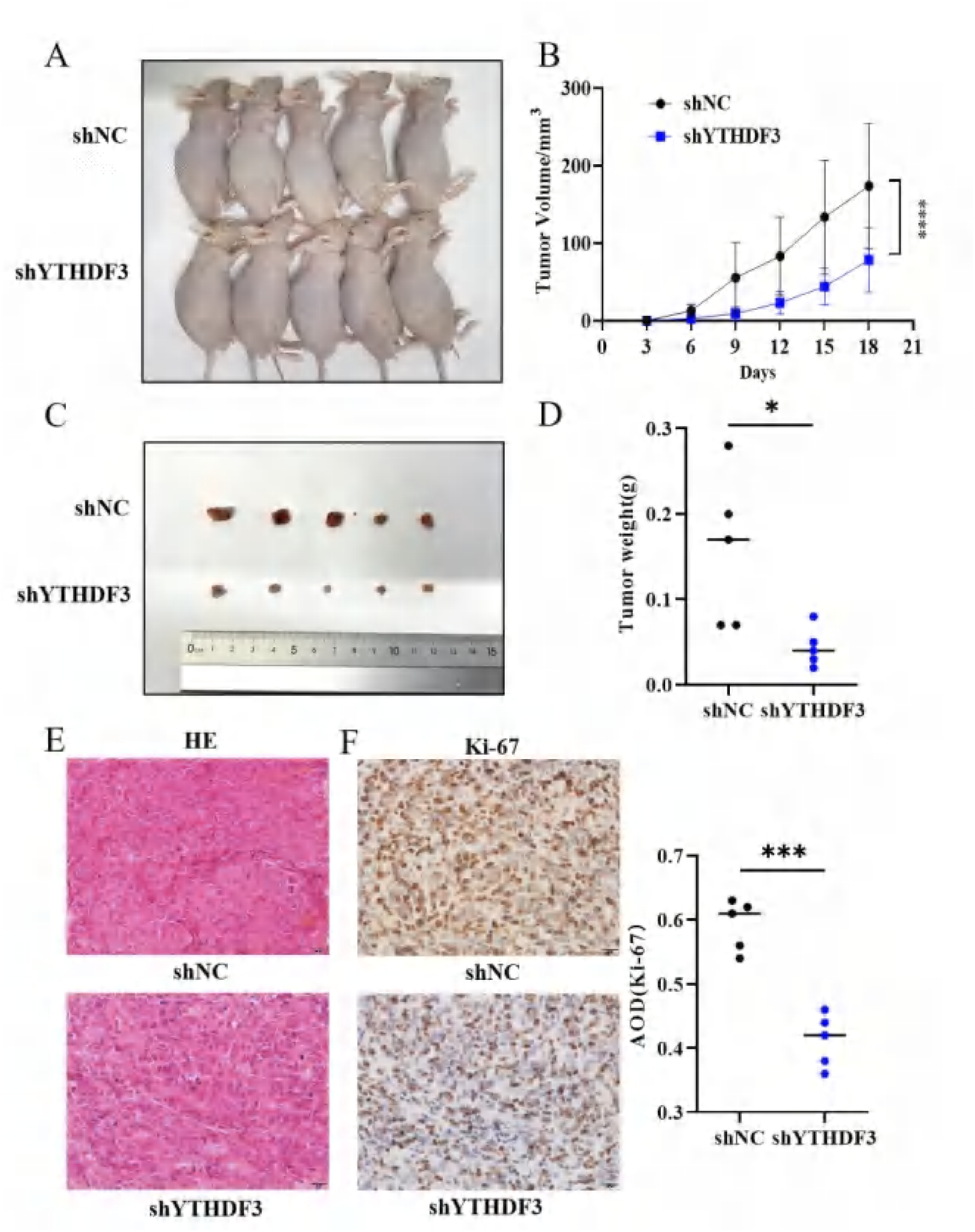
YTHDF3 regulates the growth of GC *in vivo*. (A) Nude mouse tumor-bearing model. (B) Tumor growth curve. Images (C) and weight (D) of xenografted tumors. (E) Representative images of HE-stained tumor tissue sections. Scale bar, 50 μm. (F) Representative image of Ki67 IHC staining in tumor tissue sections. Scale bar, 50 μm. Data are shown as means ± SD. \**P<*0.05, \*\*\**P<*0.001

### NEK7 was a potential target of YTHDF3 in GC

We conducted RNA-seq in HGC-27 cells of the siNC group and the siYTHDF3 group, respectively, to better investigate the possible role of YTHDF3 in the pathophysiology of GC. The RNA-seq showed that the knockdown of YTHDF3 expression significantly altered the expression of a total of 5672 genes, including up-regulated and down-regulated genes (Figure 5A, B). Through the analysis of down-regulated genes as identified by RNA-seq (fold change>2 and *P<*0.05), along with the top 1000 genes co-expressed with YTHDF3 in GC samples from the TCGA database, and the 3742 genes highly expressed in GC within the same database, two genes were identified as potential candidates for binding to the YTHDF3 protein. These two genes (NEK7/MPZL1) were potential target genes for YTHDF3 (Figure 5C). Subsequent analysis of these genes involved examining their subcellular distribution, expression in GC tissues, and the associated survival outcomes of patients. Data from TCGA database revealed that NEK7 was prominently expressed in GC tissues (Figure 5D), predominantly localized to the cytoplasm and nucleus (Figure 5E). Moreover, patients exhibiting high levels of NEK7 expression exhibited significantly reduced overall survival rates (Figure 5F). Therefore, we speculate that NEK7 is a potential downstream target gene. Co-IF assays wa<u>s</u> then used to confirm the interaction between YTHDF3 and NEK7. The Co-IF results demonstrated that YTHDF3 and NEK7 co-localized in the cytoplasm and nucleus of GC cells (Figure 5G). To determine whether YTHDF3 interacts directly with NEK7 mRNA, quantitative RNA immunoprecipitation PCR (RIP-qPCR) was performed. The results demonstrated that the YTHDF3 protein had significantly enriched NEK7 mRNA compared with that of the control IgG group (Figure 5H). This finding indicates a specific interaction between YTHDF3 and NEK7 transcripts. ActD-designed mRNA decay assays were used to block nascent transcription and revealed that YTHDF3 knockdown markedly reduced the half-life of NEK7 mRNA (Figure 5I). Lastly, we investigated the transcriptional and translational alterations of NEK7 after YTHDF3 knockdown or overexpression to further validate the regulation function of NEK7 expression by YTHDF3 in GC. After downregulating YTHDF3 expression in HGC-27 cells, both NEK7 mRNA and protein expression were reduced (Figure 5J, K). Similarly, when YTHDF3 expression was upregulated in MKN74 cells, both NEK7 mRNA and protein expression levels increased (Figures 5L, M). These events suggest that YTHDF3 regulates downstream gene expression at both the transcriptional and translational levels. In summary, these findings indicate an interaction between YTHDF3 and NEK7 in GC, and that YTHDF3 can regulate the production of the NEK7 protein. NEK7 may serve as a potential modification target of YTHDF3 in GC.

**Figure 5.**
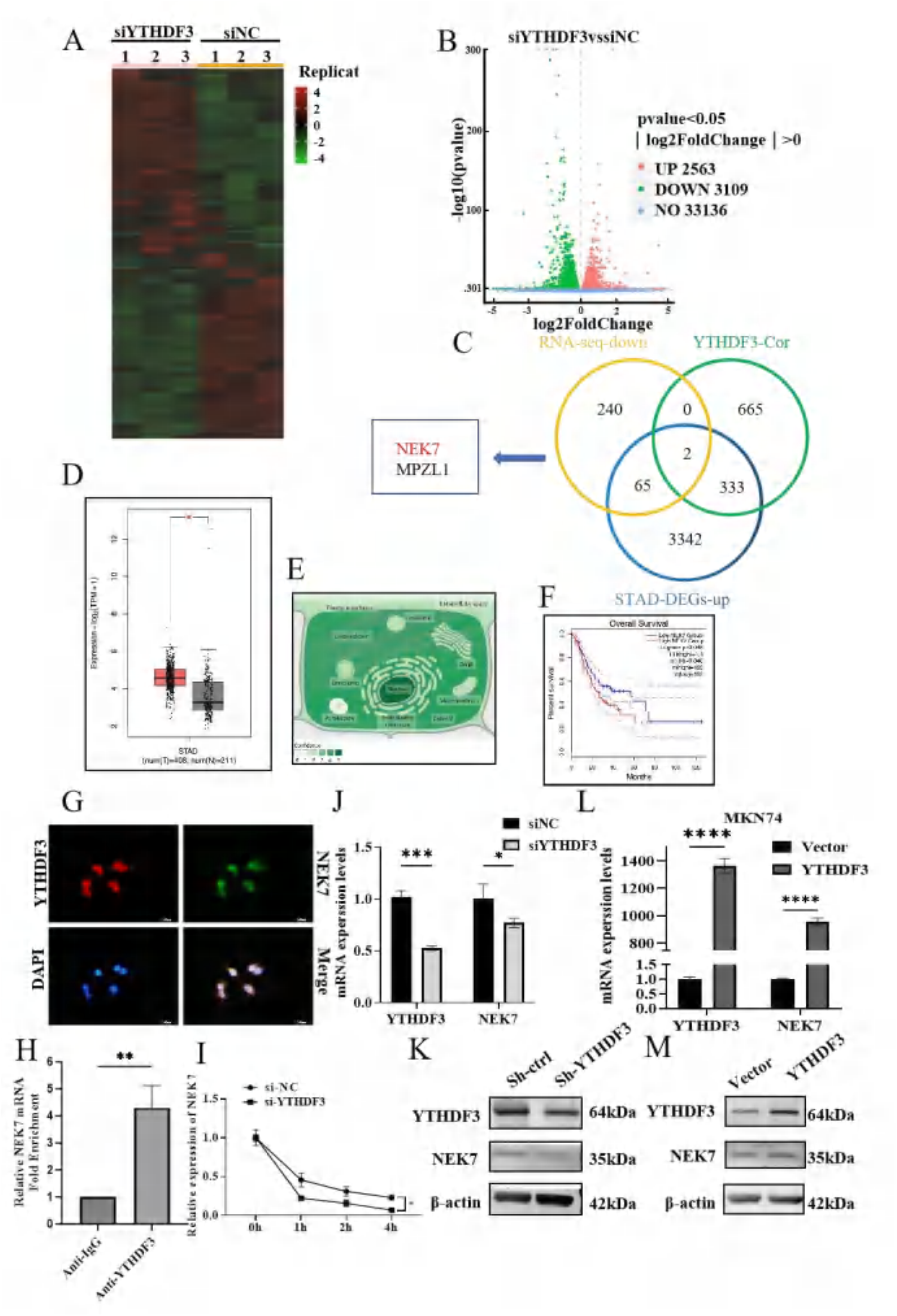
NEK7 is the modification target of YTHDF3. (A) Heatmap showing differentially expressed genes (DEGs) identified by RNA-seq. (B) Number of DEGs identified by RNA-seq. (C) Overlap analysis of genes identified with RNA-seq, YTHDF3-cor, and STAD-DEGs. (D) Expression of YTHDF3 in the GC paired tissue cohort from the TCGA database. (E) Subcellular location of NEK7 from GeneCards. (F) Survival analysis of NEK7 expression levels and overall survival in patients with GC. (G) The interaction between YTHDF3 and NEK7 as verified by the Co-IF experiments. Scale bar, 10μm. (H) The interaction between the YTHDF3 protein and the NEK7 mRNA as examined by RIP-qPCR. (I) NEK7 mRNA stability assessed by RT-qPCR after actinomycin D treatment in the siYTHDF3 and siNC groups. (J) RT-qPCR detection of NEK7 mRNA expression after YTHDF3 knockdown in HGC-27 cells. (K) Western blotting showing NEK7protein expression after YTHDF3 knockdown in HGC-27 cells. (L) RT-qPCR detection of NEK74 mRNA expression after YTHDF3 overexpression in MKN74 cells. (M) Western blotting showing NEK7 protein expression after YTHDF3 overexpression in MKN74 cells. Data are presented as means ± SD. \*\*\**P<*0.001, \*\*\*\**P<*0.0001

### NEK7 was highly expressed in GC and interacts with YTHDF3

NEK7 mRNA levels were higher than normal gastric mucosa cells (Figure 6A) and GC tissues had a higher level of NEK7 mRNA in adjacent tissues (18 GC paired tissues) (Figure 6B). MKN74 cells had the highest expression of NEK7 protein (Figure 6C), whereas GC tissues had higher levels than nearby tissues (44 GC paired tissues) (Figure 6D). These findings imply that NEK7 may play a role in GC pathogenesis as an oncogene. Subsequently, our aim was to further elucidate the mechanism by which the interaction between YTHDF3 and NEK7 contributes to the development of GC. One of the siRNA sequences with the highest knockdown efficacy (si-NEK7-659) was selected for further testing after four siRNA-NEK7 sequences were transfected concurrently into MKN74 cells with the highest expression of NEK7 (Figure 6E-F). We then transfected siRNA-negative control/siRNA-YTHDF3 and pcDNA3.1-vector/pcDNA3.1-NEK7 into HGC-27 and siRNA-negative control/siRNA-NEK7 and pcDNA3.1-vector/pcDNA3.1-YTHDF3 into MKN74 cells, simultaneously or separately. RT-qPCR and western blotting were used to identify variations in YTHDF3 expression in each group. The findings indicated that overexpression of NEK7 mitigated the effects of YTHDF3 knockdown in HGC-27 cells (Figure 6J-H). Furthermore, the knockdown of NEK7 also mitigated the effects of YTHDF3 upregulation in MKN74 cells (Figure 6I, J). These results suggest that NEK7 can influence the expression of YTHDF3 by GC cells.

**Figure 6.**
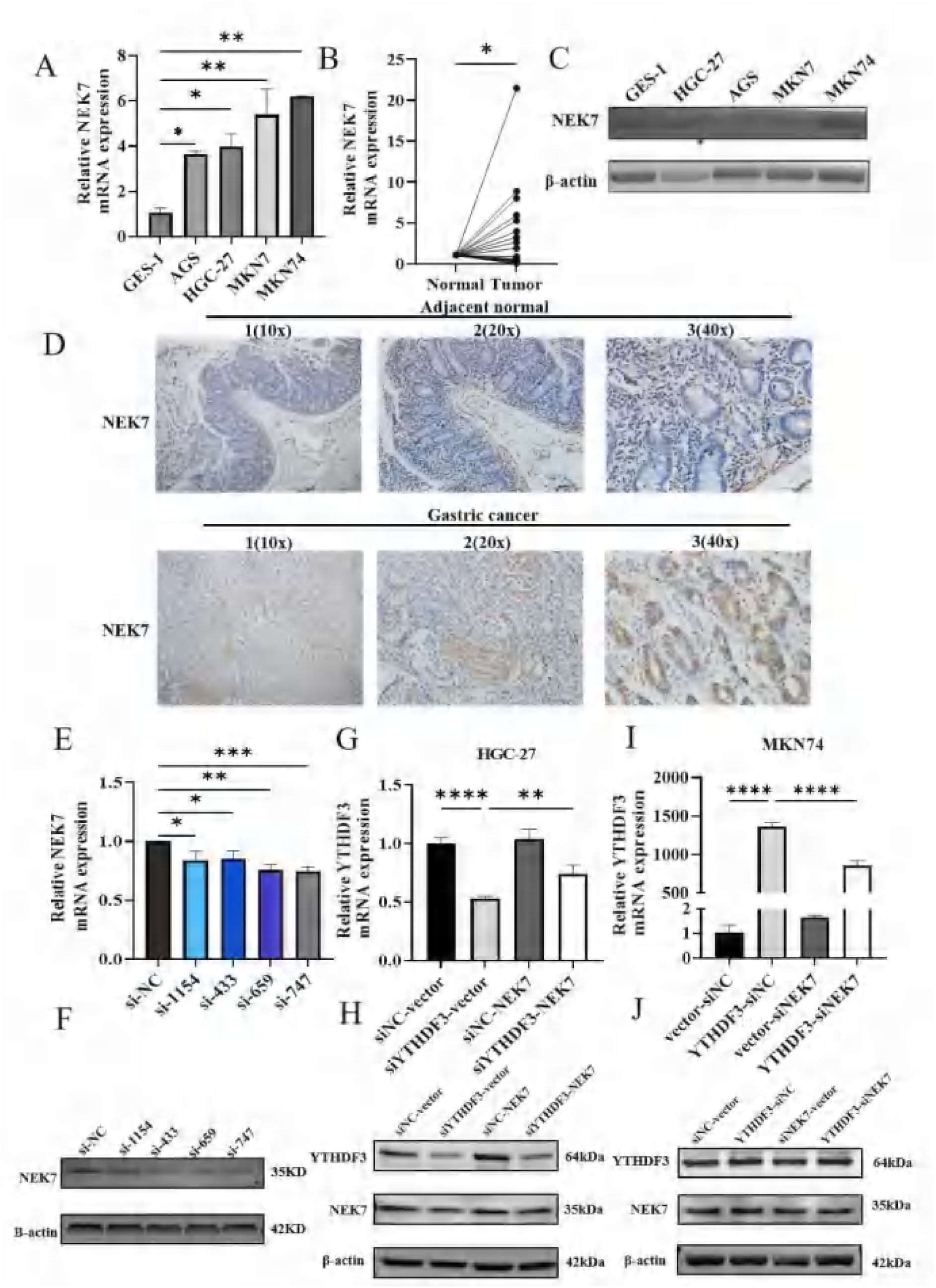
Expression of NEK7 and its interaction with YTHDF3 in GC. (A) NEK7 mRNA expression in GC cell lines and GES-1 as detected by RT-qPCR. (B) NEK7 mRNA expression in 18 pairs of GC paired tissues was detected by RT-qPCR. (C) The basal expression of NEK7 protein in cell lines as detected by WB. (D) NEK7 protein expression in 44 pairs of GC paired tissues as detected by IHC. Scale bar, 50 μm. (E-F) Four siRNA-NEK7 sequences were transfected simultaneously into MKN74 cells. (G-H) The relative expression of YTHDF3 mRNA and protein as detected by RT-qPCR and western blotting, respectively, after instantaneous co-transfection of YTHDF3 siRNA and NEK7 overexpression plasmid in HGC-27. (I-J) The relative expression of YTHDF3 mRNA and protein were detected by RT-qPCR and western blotting, respectively, after instantaneous co-transfection of NEK7 siRNA and YTHDF3 overexpression plasmids in MKN74. Data are presented as means ± SD. \*\*\**P<*0.001, \*\*\*\**P<*0.0001

### YTHDF3 targeted NEK7 to activate the Wnt/β-catenin signaling pathway in cancer promotion

We performed KEGG pathway enrichment analysis of the DEGs identified using RNA-seq data and examined the signaling pathways regulated by them to elucidate the oncogenic mechanism of YTHDF3 in GC. The findings indicated a high enrichment of genes in the Wnt/β-catenin signaling pathway (Figure 7A). We then investigated whether NEK7 could reverse the inhibitory effects of the suppression of YTHDF on the activation of Wnt/β-catenin signaling. The findings imply that downstream effectors of β-catenin (c-MYC, CD1) had similarly markedly decreased expression in HGC-27 cells on downregulation of YTHDF3 expression. At the same time, overexpression of NEK7 can reverse the downregulation of these effectors caused by the loss of YTHDF3 (Figure 7B). In contrast, YTHDF3 overexpression increased the levels of downstream effectors of β-catenin (c-MYC and CD1) in HGC-27 cells, and this up-regulation was abolished by concurrent suppression of NEK7 (Fig. 7C). Immunofluorescence staining showed a significantly stronger nuclear fluorescence intensity in HGC-27 cells transfected with pcDNA3.1-NEK7 compared with the pcDNA3.1-NC controls (Fig. 7D), indicating that NEK7 over-expression promotes nuclear translocation of β-catenin in GC cells. Western blotting analysis revealed that NEK7 overexpression significantly reduced total phosphorylated-β-catenin protein levels in HGC-27 cells (Fig. 7E). These results imply that YTHDF3 regulates the activation of the Wnt/β-catenin signaling pathway via NEK7. We then investigated whether NEK7 expression had an functional role on YTHDF3 activity in GC cells. The upregulation of NEK7 expression restored the attenuation of cell proliferation, migration, and invasion in HGC-27 cells caused by YTHDF3 knockdown (Figure 7F,G,J,L,M). The downregulation of NEK7 expression mitigated the augmentation of cell proliferation, migration, and invasion induced by YTHDF3 overexpression in MKN74 cells (Figure 7H,I,K,N,O). Altogether, the findings indicate that NEK7 is an essential downstream target for YTHDF3 that aids in the development of GC, and NEK7 and the Wnt/β-catenin signaling pathway are necessary for YTHDF3 to have an oncogenic effect on GC. Thus, we suggest that higher expression of YTHDF3 may enhance the production of NEK7 mRNA and protein, which in turn may activate the Wnt/β-catenin signaling pathway and eventually favor tumors development of GC cells. To further investigate the involvement of the Wnt/β-catenin pathway in YTHDF3-mediated oncogenesis, we treated gastric adenocarcinoma cells with IWR-1-endo, a specific Wnt/β-catenin pathway inhibitor. HGC-27 cells with high endogenous expression of YTHDF3 were divided into four treatment groups: si-NC+DMSO, si-YTHDF3+DMSO, si-NC+IWR-1-endo, and si-YTHDF3+IWR-1-endo. We first determined the half-maximal inhibitory concentration (IC50) of IWR-1-endo in HGC-27 cells using the CCK-8 assay (Fig. 7P. Subsequently, cells were treated with IWR-1-endo at the determined concentration for functional analyses. The CCK-8 assay revealed that the knockdown of YTHDF3 significantly enhanced the suppressive effects of IWR-1-endo on cell proliferation (Fig. 7Q). Collectively, these results demonstrate that the oncogenic functions of YTHDF3 are mediated, at least in part, through the Wnt/β-catenin signaling pathway.

**Figure 7.**
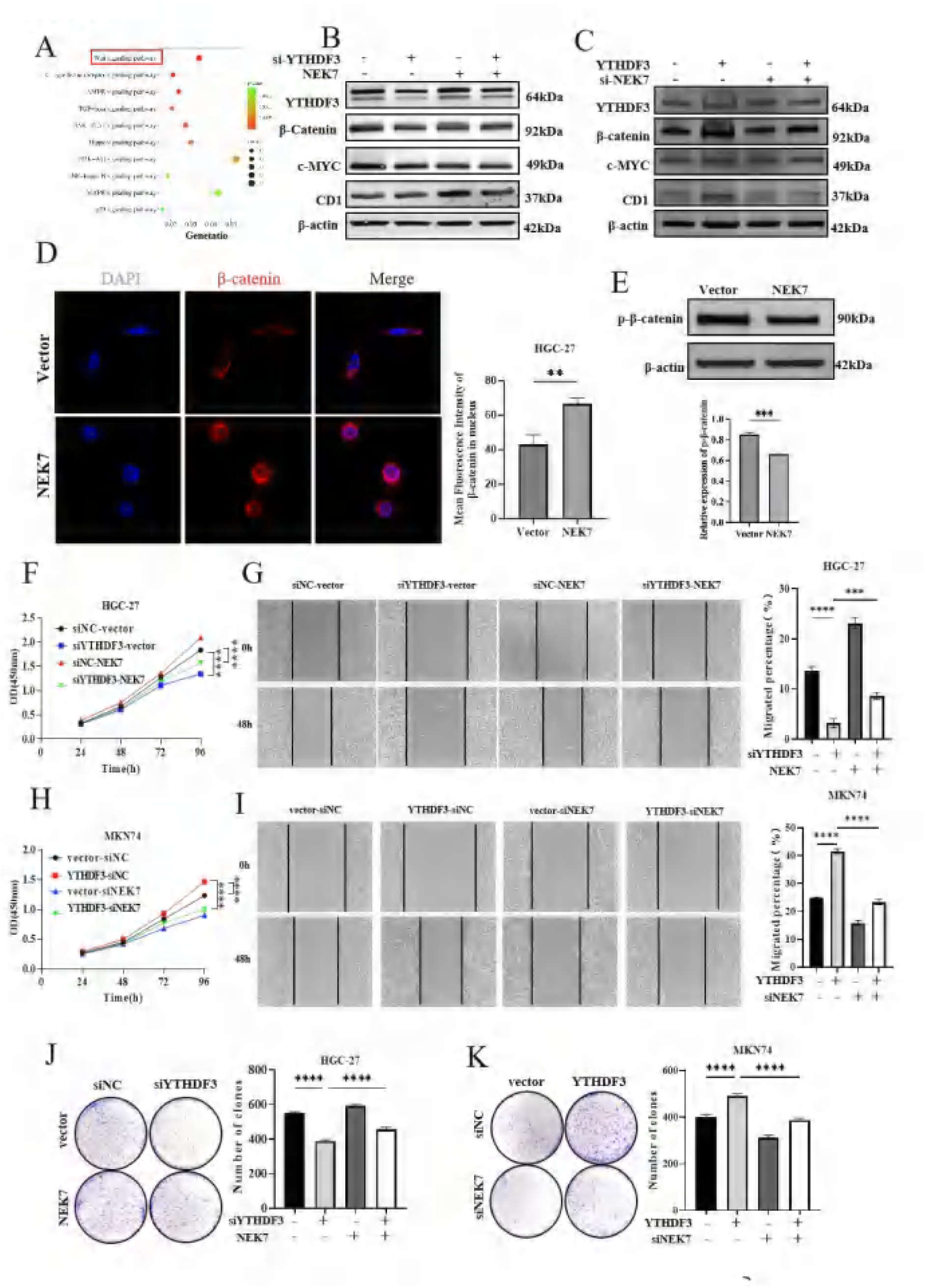
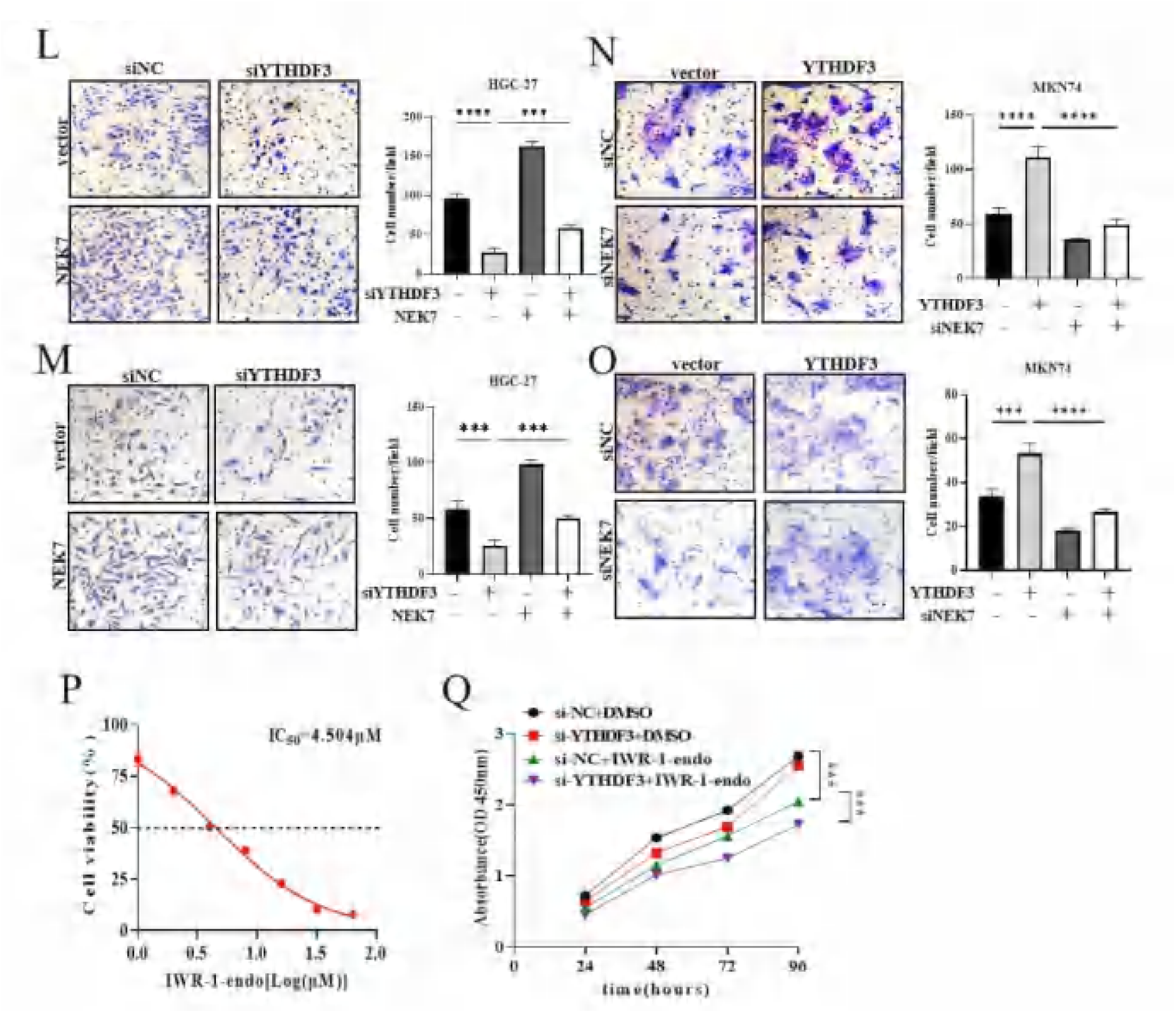
Oncogenic function of YTHDF3 depends on NEK7 and Wnt/β-catenin signaling pathway. (A) KEGG enrichment analysis of differentially expressed gene sets from the RNA-seq data. (B) The expression of Wnt/β-catenin signaling pathway related proteins as detected by western blotting following instantaneous co-transfection of YTHDF3 siRNA and NEK7 overexpression plasmid in HGC-27. (C) Expression of Wnt/β-catenin signaling pathway related proteins as detected by western blotting after instantaneous co-transfection of YTHDF3 overexpression and NEK7 siRNA plasmid in HGC-27. (D) Nuclear accumulation of β-catenin in HGC-27 cells over-expressing NEK7 analyzed by immunofluorescence. (E) Western blotting determination of total phosphorylated-β-catenin protein levels in NEK7-overexpressing HGC-27 cells. The effects of NEK7 on YTHDF3 in HGC-27 cells were detected by the CCK-8 assay (F), wound healing assay (G), Scale bar, 200 μm, colony formation assay (J), Transwell migration assay (L) and Transwell invasion assay (M). Scale bar, 100 μm. Effects of NEK7 on YTHDF3 in MKN74 cells as detected by t h e CCK-8 assay (H), wound healing assay (I), Scale bar, 200 μm, colony formation assay (K), Transwell migration assay (N), and Transwell invasion assay (O). Scale bar, 100 μm. (P) The IC50 of WR-1-endo in HGC-27 cells was determined by the CCK-8 assay. (Q) Cell proliferation was assessed by the CCK-8 assay in HGC-27 cells treated with 4.504 μM WR-1-endo. Data are presented as means ± SD. \**P<*0.05, \*\**P<*0.01, \*\*\**P<*0.001, \*\*\*\**P<*0.0001

## Discussion

This study reveals that YTHDF3 expression is significantly upregulated in patients with GC and is closely associated with a poor prognosis. Using both *in vitro* and *in vivo* models, YTHDF3 was shown to exhibit characteristics of an oncogene. Knockdown of YTHDF3 expression markedly inhibited the proliferation, migration, and invasion of GC cells while promoting apoptosis, whereas overexpression of YTHDF3 had the opposite effect. Further, the study showed that YTHDF3 exerted its oncogenic role by targeting NEK7 and activating the Wnt/β-catenin signaling pathway, which suggests that YTHDF3 may promote tumor development in GC cells by improving the production of NEK7 mRNA and protein.

Previous studies have found that YTHDF3 expression was significantly higher in HCC tissues than in paracancerous liver tissues. YTHDF3 was also significantly upregulated in HCC with microvascular invasion (MVI) compared with HCC without MVI (19). This is consistent with our findings of elevated levels of expression of YTHDF3 mRNA and proteins in GC tissues. Further analysis of the relationship between YTHDF3 expression and clinical pathological data in patients with GC revealed that high YTHDF3 expression was closely associated with lymph node metastasis and more advanced TNM stages, which holds significant value for individualized clinical treatment and prognostic evaluation.

To validate the impact of YTHDF3 on the biological functions of GC cells, our *in vitro* experiments demonstrated that knockdown of YTHDF3 in GC cells significantly suppressed cell proliferation, migration, and invasion, and increased apoptosis. In contrast, upregulation of YTHDF3 in GC cells produced the opposite effects. Moreover, *in vivo* experiments in nude mice showed that YTHDF3 depletion inhibited the growth of gastric tumors. Overexpression of YTHDF3 has also been shown to facilitate the proliferation, invasion, and migration of HCC cells both *in vitro* and *in vivo*; however, YTHDF3 knockdown resulted in an inverse trend (17). YTHDF3 is required for the maintenance of CSC properties and tumor initiation capacity in ocular melanoma *in vitro* and *in vivo*. Ocular melanoma cells with targeted YTHDF3 knockdown exhibited inhibitory tumor proliferation and migration abilities (13). This aligns with our findings, collectively confirming that YTHDF3 can facilitate malignant biological characteristics such as growth and metastasis, potentially contributing to the development of GC.

Subsequently, we sought to identify the target genes that mediate the effects of YTHDF3 to elucidate the molecular mechanisms of GC. Through RNA-seq, downregulated gene analysis, YTHDF3 correlation analysis, and STAD-DEGs-up gene overlap analysis, combined with online database analysis and literature review, NEK7 drew our attention. We confirmed its binding to YTHDF3 by Co-IF and RIP-qPCR followed, which confirmed the interaction between YTHDF3 and NEK7 in GC cells, a finding not previously reported. NEK7 is a multifunctional serine/threonine kinase expressed in various organs and is crucial for cell division, mitochondrial function, and protein transport. Studies have shown that NEK7 is highly expressed in multiple cancers and is closely associated with tumor growth, invasion, and metastasis (19). In particular, the role of NEK7 has been confirmed in retinoblastoma, breast cancer, hepatocellular carcinoma, and pancreatic cancer (20–22). Fortunately, these results are consistent with our current findings on NEK7 in GC. To explore the mechanism by which YTHDF3 regulates NEK7 in GC, we demonstrated that YTHDF3 regulates NEK7 expression at the translational and transcriptional levels. We determined that YTHDF3 can bind to NEK7 mRNA and promote its translation, ultimately leading to the progression of GC.

Enrichment analysis of DEGs from RNA-seq in the KEGG pathway analysis revealed significant enrichment in the Wnt/β-catenin signaling pathway. The classic WNT/β-catenin signaling pathway maintains high stability during evolution and plays a key regulatory role in tissue formation and homeostasis (23). This pathway plays a central role in cancer formation, progression, and the immune response to therapy. Under normal physiological conditions, it is responsible for regulating cell growth, differentiation, and programmed cell death; however, in cancer cells, its overactivation typically promotes tumor growth and spread and helps tumor cells evade immune surveillance. Numerous studies have shown that Wnt signaling pathway-related proteins are widely distributed in various types of cancer, such as colorectal cancer (24), lung cancer (25), malignant melanoma (26), breast cancer (27), hepatocellular carcinoma (28), and pancreatic cancer (29). These proteins are located in membrane structures, cytoplasm, and nuclei of tumor cells. Clearly, the Wnt/β-catenin pathway plays a dominant role in the initiation and evolution of cancer, driving the occurrence and malignant transformation of cancer cells (30,31). Based on the analysis presented above, we hypothesized that high expression of YTHDF3 may promote the oncogenic effects of the Wnt/β-catenin signaling pathway. Subsequently, we used western blotting to detect key markers of the Wnt/β-catenin pathway, including β-catenin, c-MYC, and CD1. Furthermore, the inhibitory effects of YTHDF3 knockdown on Wnt/β-catenin pathway activation could be rescued by NEK7 overexpression, and NEK7 itself was capable of activating the Wnt/β-catenin signaling pathway. In subsequent rescue experiments, we further confirmed that the YTHDF3/NEC7/Wnt/β-catenin axis influenced the growth, migration, and invasion capabilities of GC cells. Overall, these results suggest that the YTHDF3/NEF7/Wnt/β-catenin axis is involved in the occurrence and development of GC, providing potential molecular targets for the diagnosis and treatment of GC, and laying the foundation for further research on the role of the YTHDF family in cancer development.

Mechanistically, our study demonstrates that YTHDF3 promotes GC progression by binding and stabilizing NEK7 mRNA, thus activating the Wnt/β-catenin pathway. This mechanism underscores the functional complexity of YTHDF3 as an m6A reader. Unlike YTHDF1, which primarily promotes mRNA translation, and YTHDF2, which mainly mediates mRNA decay, YTHDF3 possesses a more flexible dual regulatory capacity. YTHDF2 can facilitate the translation of target mRNA in concert with YTHDF1, while promoting mRNA decay in cooperation with YTHDF2 (32). In our study, YTHDF3 mainly acted to stabilize NEK7 mRNAand further enhanced its translation, which is in contrast with the canonical function of YTHDF2, but is aligned with reports positioning YTHDF3 as a stabilizing factor in certain contexts (33). This suggests that the YTHDF proteins do not function in strict isolation but rather form a dynamic and context-dependent network regulating RNA metabolism.

Our findings provide new evidence for the cancer-specific functions of YTHDF3. For example, in NSCLC tumor, YTHDF3 drives progression by promoting the decay of tumor suppressor mRNA (34), which contrasts strongly with the mRNA stabilization mechanism observed in our study. Similarly, in colorectal cancer, YTHDF3 operates by directly enhancing the translation efficiency of oncogenic transcripts (35). These discrepancies highlight the highly cancer-type-dependent nature of YTHDF3 regulation. Our study is the first to establish a direct link of YTHDF3 to the Wnt/β-catenin pathway in GC. Previous studies have demonstrated that the Wnt/β-catenin pathway is pivotal in GC, its upstream regulation by YTHDF3, particularly through the NEK7 node, has not been previously reported. Therefore, our study not only reveals a novel mechanism of action for YTHDF3 in GC but also suggests that targeting the YTHDF3-NEK7-Wnt/β-catenin axis could represent a potential new therapeutic strategy.

In conclusion, this study identified YTHDF3 as a key promoter of GC progression through stabilization of NEK7 and activation of the Wnt/β-catenin. Our findings provide a rationale for considering the YTHDF3/ NEK7 axis as a potential therapeutic target for the treatment of GC. The development of specific YTHDF3 inhibitors and their subsequent testing in pre-clinical models represent critical next steps. Ultimately, validating YTHDF3-NEK7-Wnt/β-catenin axis in clinical settings will be essential for translating our mechanistic findings into future therapeutic strategies for patients with GC.

## Availability of data and materials

The datasets generated and/or analyzed during the current study are available from the corresponding author on reasonable request. The materials used in this study are commercially available or described in the Methods section.

## Authors’ contributions

Conceptualization and experimentation: XRS, YFC, TL. Administrative support: XRS, WFY, SSH. Resource provision: RZ, JJ. Data collection and curation: MDX, KYL. Data analysis and interpretation: JW, YS, XRS. All authors participated in manuscript writing and final approval.

## Ethics approval and consent to participate

The research involving human subjects was conducted in accordance with the principles of the Helsinki Declaration and the International Ethical Guidelines for Biomedical Research Involving Human Subjects, and was approved by the Ethics Committee of the First Hospital of Hebei Medical University (Approval No. S00996). The animal experiments were carried out in compliance with the Guiding Opinions on the Kind Treatment of Experimental Animals, the Guidelines for the Ethical Review of Animal Welfare in Experimental Animals, and the Guiding Principles for the Registration Review of Medical Device Animal Trials, and were authorized by the Ethics Committee of the First Hospital of Hebei Medical University (Approval No. S00994).

## Patient consent for publication

Not applicable

## Abbreviations

ANOVA: Analysis of variance
AOD: Average optical density
CT: Cyclic threshold
DEG: Differentially expressed genes
FBS: Fetal bovine serum
GC: Gastric cancer
GEO: Gene Expression Omnibus
RT: Reverse-transcribe
STAD: Stomach adenocarcinoma
TCGA: The Cancer Genome Atlas
YTHDF: YTH domain family

## References

1. Bray F, Ferlay J, Soerjomataram I, et al. Global cancer statistics 2022: GLOBOCAN estimates of incidence and mortality worldwide for 36 cancers in 185 countries. CA Cancer J Clin. 2024 May-Jun;74(3):229–263.

2. Zaccara S, Patil DP, Jaffrey SR. A unified model for the function of YTHDF proteins in regulating m6A-modified mRNA. Cell. 2020;181:1582–1595.

3. Li L, Tang C, Ye J, et al. Bioinformatic analysis of m6A “reader” YTH family in pan-cancer as a clinical prognosis biomarker. Sci Rep. 2023;13:17350.

4. Mei Z, Shen Z, Pu J, et al. NAT10 mediated ac4C acetylation driven m6A modification via involvement of YTHDC1-LDHA/PFKM regulates glycolysis and promotes osteosarcoma. Cell Commun Signal. 2024;22:51.

5. Liao Y, Liu Y, Yu C, et al. HSP90β impedes STUB1-induced ubiquitination of YTHDF2 to drive sorafenib resistance in hepatocellular carcinoma. Adv Sci. 2023;10:e2302025.

6. Song JK, You GH, Yin XH, et al. Overexpression of YTHDC2 contributes to the progression of prostate cancer and predicts poor outcomes in patients with prostate cancer. J Biochem Mol Toxicol. 2023;37:e23308.

7. Chen J, Sun Y, Xu X, et al. YTH domain family 2 orchestrates epithelial-mesenchymal transition/ proliferation dichotomy in pancreatic cancer cells. Cell Cycle. 2017;16:2259–2271.

8. Song P, Li X, Chen S, et al. YTHDF1 mediates N-methyl-N-nitrosourea-induced gastric carcinogenesis by controlling HSPH1 translation. Cell Prolif. 2024;57:e13619.

9. Sheng H, Li Z, Su SX, et al. YTH domain family 2 promotes lung cancer cell growth by facilitating 6-phosphogluconate dehydrogenase mRNA translation. Carcinogenesis. 2020;41(5):541–550.

10. Li J, Ahmad M, Sang L, et al. O-GlcNAcylation promotes the cytosolic localization of the m6A reader YTHDF1 and colorectal cancer tumorigenesis. J Biol Chem. 2023;299:104738.

11. Ries RJ, Zaccara S, Klein P, et al. m6A enhances the phase separation potential of mRNA. Nature. 2019;571:424–428.

12. Lin Y, Chen X, Huang Y, et al. YTHDF3 facilitates triple-negative breast cancer progression and metastasis by stabilizing ZEB1 mRNA in an m6A-dependent manner. Ann Transl Med. 2022 Jan;10(2):83.

13. Xu Y, He X, Wang S. The m6A reading protein YTHDF3 potentiates tumorigenicity of cancer stem-like cells in ocular melanoma through facilitating CTNNB1 translation. Oncogene. 2022;41(9):1281–1297.

14. Liu D, Li Z, Zhang K, et al. N-methyladenosine reader YTHDF3 contributes to the aerobic glycolysis of osteosarcoma through stabilizing PGK1 stability. J Cancer Res Clin Oncol.

15. Ni W, Yao S, Zhou Y, et al. Long noncoding rna Gas5 inhibits progression of colorectal cancer by interacting with and triggering yap phosphorylation and degradation and is negatively regulated by the M6A reader Ythdf3. Mol Cancer. 2019;18(1):143.

16. Hu Y, Zhang Y, Chen X, et al. A reciprocal feedback between N6-methyladenosine reader YTHDF3 and lncRNA DICER1-AS1 promotes glycolysis of pancreatic cancer through inhibiting maturation of miR-5586-5p. J EXP Clin Cancer Res. 2022 Feb 19;41(1):69.

17. Hu B, Gao J, et al. m6A reader YTHDF3 triggers the progression of hepatocellular carcinoma through the YTHDF3/m6A-EGFR/STAT3 axis and EMT. Mol Carcinog. 2023 Jul 14;62(10):1599–1614.

18. Zhao Y, Zhang Y, Li Y, et al. YTHDF3 Facilitates eIF2AK2 and eIF3A Recruitment on mRNAs to Regulate Translational Processes in Oxaliplatin-Resistant Colorectal Cancer. ACS Chemical Biology. 2022 Jul 15;17(7):1778–1788.

19. Eisa NH, Jilani Y, Kainth K, et al. The co-chaperone UNC45A is essential for the expression of mitotic kinase NEK7 and tumorigenesis. J Biol Chem. 2019;294:5246–5260.

20. Kooi IE, Mol BM, Massink MP, et al. A Meta-Analysis of Retinoblastoma Copy Numbers Refines the List of Possible Driver Genes Involved in Tumor Progression. PLoS One. 2016;11(4):e0153323.

21. Zhang J, Wang L, Zhang Y. Downregulation of NIMA-related kinase-7 inhibits cell proliferation by inducing cell cycle arrest in human retinoblastoma cells. Exp Ther Med. 2018;15(2):1360–1366.

22. Yan Z, Qu J, Li Z, et al. NEK7 Promotes Pancreatic Cancer Progression And Its Expression Is Correlated With Poor Prognosis. Front Oncol. 2021;11:705797.

23. Liu D, Chen L, Zhao H, et al. Small molecules from natural products targeting the Wnt/beta-catenin pathway as a therapeutic strategy. Biomed Pharmacother. 2019;117:108990.

24. O’Brien CA, Pollett A, Gallinger S, et al. A human colon cancer cell capable of initiating tumour growth in immunodeficient mice. Nature. 2007;445(7123):106–10.

25. Pan JC, Fang S, Tian HH. lncRNA JPX/miR-33a-5p/Twist1 axis regulates tumorigenesis and metastasis of lung cancer by activating Wnt/β-catenin signaling. Mol Cancer. 2020;19(1):9.

26. Dong F, Nguyen T, Kim L, et al. A tumorigenic subpopulation with stem cell properties in melanomas. Cancer Res. 2005;65(20).

27. Chen ZH, Tian Y. CMTM7 inhibits breast cancer progression by regulating Wnt/β-catenin signaling. Breast Cancer Res. 2023;25(1):22.

28. Ma S, Chan KW, Hu L, et al. Identification and characterization of tumorigenic liver cancer stem/progenitor cells. Gastroenterology. 2007;132(7):2542–56.

29. He JL, Yang GH, Cao RL, et al. RGC-32 facilitates pancreatic cancer via activating Wnt/β-catenin signaling. Cell Mol Biol. 2023;69(14):161–165.

30. Verras M, Sun Z. Roles and regulation of Wnt signaling and beta-catenin in prostate cancer. Cancer Lett. 2006;237(1).

31. Wend P, Holland JD, Ziebold U, et al. Wnt signaling in stem and cancer stem cells. Semin Cell Dev Biol. 2010;21(8).

32. Shi H, Wang X, Lu Z, et al. YTHDF3 facilitates translation and decay of N6-methyladenosine-modified RNA. Cell Research, 2017;27(3):315–328.

33. Hu B, Gao J, Shi J, Wen P, et al. m A reader YTHDF3 triggers the progression of hepatocellular carcinoma through the YTHDF3/m A-EGFR/STAT3 axis and EMT. Mol Carcinog. 2023;62(10):1599–1614.

34. Jin D, Guo J, Wu Y, et al. mA demethylase ALKBH5 inhibits tumor growth and metastasis by reducing YTHDFs-mediated YAP expression and inhibiting miR-107/LATS2-mediated YAP activity in NSCLC. Mol Cancer. 2020 Feb 27;19(1):40.

35. Zeng K, Peng J, Xing Y, et al. A positive feedback circuit driven by mA-modified circular RNA facilitates colorectal cancer liver metastasis. 2023;22(1):202.

